# Decoupling Genetics from Attainments: The Role of Social Environments

**DOI:** 10.1101/817163

**Authors:** Jason Fletcher

## Abstract

This paper examines the extent to which growing up in a socially mobile environment might decouple genetic endowments related to educational attainment with actual attainments. Many models of intergenerational transmission of advantage contain both a transmission channel through endowments (i.e. genetics) from parents to children as well as from parental investments and “luck”. Indeed, many scholars consider the intergenerational links due to the transmission of genetically based advantage to place a lower bound on plausible levels of social mobility—genetics may be able to “lock in” advantage across generations. This paper explores this idea by using new genetic measurements in the Health and Retirement Study to examine potential interactions between social environments and genetics related to attainments. The results suggest evidence of gene environment interactions: children born in high mobility states have lower genetic penetrance—the interaction between state-level mobility and the polygenic score for education is negative. These results suggest a need to incorporate gene-environment interactions in models of attainment and mobility and to pursue the mechanisms behind the interactions.

## Introduction

Large literatures across several social science disciplines have been concerned with examining the magnitude and mechanisms of the intergenerational transmission of socioeconomic status (Black and Devereux 2010). Key components of the process are often separated into an endowment pathway and an investments/luck pathway or “nature” and “nurture”. Within the endowment pathway, the core focus has been on genetics as a typically unobserved but important reason that the outcomes of children and their parents are positively correlated. To a large extent, understanding the importance of non-genetic pathways is challenging both because genetics is thought to operate across a wide range of domains and because it is often unobserved by the researcher.^2^ Emerging work has begun to challenge the traditional nature vs. nurture decomposition exercise to instead consider the interplay of nature and nurture (Heckman 2007, 2008), or gene-environment interactions. However, much evidence has not been able to separate gene-environment interactions from gene-environmental correlations (Fletcher and Conley 2013).^3^ For analyses of attainments, gene-environment correlation occurs in part due to the positive relationship between children’s genetics, parents’ genetics, and parentally provided environments (Belsky et al. 2018).

This paper contributes to the literature on how genetics, environments, and their interaction are related to adult attainments in several ways. First, following a small but growing set of papers, this paper incorporates new genetic measurements to the study of socioeconomic attainment (Belsky et al. 2018, Conley et al. 2015). In particular, I use a summary measure of genetic endowments, a polygenic score^4^ for education developed by a massive Genome Wide Association Study (GWAS) by Okbay et al. (2016). Second, I contribute a new measure of social environments by estimating a measure of education mobility at the state-birth decade level by leveraging the Health and Retirement Study. Combining these two measures then allows the key contribution of the paper: a novel examination of whether genetics are decoupled from attainments to a larger extent in socially mobile environments (i.e a gene-environment interaction).^5^ The main results suggest evidence of gene environment interactions. Indeed, children growing up in high mobility states have lower genetic penetrance—the interaction between state-level mobility and the polygenic score for education is negative.

### Background Literature

Deep literatures across several social sciences, including sociology, demography, and economics, have focused attention on theoretic and empirical models predicting socioeconomic attainments in adulthood. There are volumes of evidence on the variation in of these predictors, including the role of parents and social institutions such as schools. Especially in analyses that explore questions of social mobility—the extent to which children’s adult attainments are tied to their parents’ attainments—a fundamental question is the extent to which genetic transmission of parental traits “lock in” some level of persistence in attainments, or immobility. Missing from this question and the typical modeling of transmission is the possibility that environments can act on and interact with genetics, which could vary levels of genetic penetrance and therefore “unlock” persistence in attainments.

While the importance of gene-environment interaction in determining socioeconomic status attainments is conceptually attractive, providing direct empirical evidence of interaction presents many challenges. Most studies are unable to leverage research designs that can separate gene-environment interaction effects from other processes. In particular, because parents contribute to both the genetic and environmental advantages/disadvantages of their children (labeled gene-environment correlation in the literature), it is difficult to separately estimate the main effects of genetics, environments, and their interactions. This is an important issue, as Belsky et al. (2016, 2018) have shown that advantageous genetic and environment factors are positively correlated. Much of this work has not been able to measure genotype explicitly, relying instead on family based studies and comparisons of heritability estimates (genes) across measures of environment, such as parental socioeconomic status^6^.

With newly available measures of genotype available in large social science surveys, a small number of studies have examined gene-environmental effects related to determinants of socioeconomic status, and very few have been able to use research designs aimed at uncovering causal estimates^7^. Conley et al. (2015) estimate associations between parental education and child genotype using the Framingham Heart Study and Health and Retirement Study and find no evidence of interaction, though this analysis was not able to separate gene-environment interaction from gene-environment correlation^8^. Thompson (2014) examines interactions between family income and a specific variant on the *MAOA* gene in predicting educational attainments of children in the Add Health data; he is able to use sibling differences in the genetic variant (a so-called “genetic lottery”^9^) to break the correlation between genetic and environment factors. The author found that children with a particular variant of *MAOA* seem to be relatively unaffected by family income variation^10^, though using a sibling difference model assumes no spillover between siblings (Boardman and Fletcher 2015).

Papageorge and Thom (2016) also use the HRS data and estimate associations between childhood socioeconomic status indicators and a polygenic score for education in determining a range of adult outcomes (education, wages, etc). While the findings suggest interactions, the research design is unable to fully separate gene-environment interaction from gene-environment correlation, since the measures of childhood socioeconomic status are largely determined by their parents (and their parents’ genotypes).

This study extends the literature by focusing attention on measures of the environment that are more plausibly not directly determined by children’s/parent’ genotype. At the same time, the state-level measure that is created and described below provides a useful omnibus test of the hypothesis that mobile environments can influence levels of genetic penetrance and therefore is an important component of understanding variation in levels of social mobility. This study also merges the literature on gene-environment interactions to explore larger scale measures of the environment by focusing on state-of-birth measures of educational mobility.

### Data

This paper uses the Health and Retirement Study (HRS), a nationally representative, longitudinal panel study of individuals over the age of 50 and their spouses. The HRS introduces a new cohort of participants every six years and interviews around 20,000 participants every two years. While the HRS has collected data on over 10,000 respondents beginning in 1992 (and refreshed in ongoing surveys), genetic data (from saliva samples) were first collected in 2006.^11^ Together with additional collections in 2008 and 2010, the genetic subsample of HRS now has over 12,000 subjects. The genetic data is comprised of over 2.5 million genetic locations for each respondent, of the over 3 billion locations in the human genome^12^. The data reports, at each location, whether the respondent has an A, C, G, or T nucleotide. The 2.5 million locations were chosen to focus on places in the genome that differ in humans, “common variants”, in at least 1% of the human populations and measure single nucleotide polymorphisms (SNP).^13^

The 2.5 million locations in the HRS are then used to create a polygenic score for education attainment. Following the literature (Belsky et al. 2016, Papageorge and Thom 2016, etc.), this score is created by weighing each nucleotide by the estimated beta coefficient linking each location with educational attainment from a massive Genome Wide Association Study (GWAS) by Okbay et al. (2016). This polygenic score captures 7% of the variation in reported educational attainment in the HRS data.

The key hypothesis in this paper is that socially mobile environments reduce the genetic penetrance for educational attainment. There is no single-way to measure “socially mobile environments.” Using a measure at a highly aggregated level (i.e. higher than individual-, family-, school-, neighborhood-level) allows the advantage of reducing the likelihood of parents choosing the environment for their children, which would induce gene-environment correlation. A limitation of a highly aggregated measure is that, while it can provide an omnibus test of the hypothesis, it cannot otherwise ascertain the particular components of the environment reasonable for the resulting decoupling of genetics from attainments.

A second issue with assessing aggregated measures, is that there are no available contextual measures of social mobility available at sub-national (e.g. state) levels in the US for the first half of the 20^th^ century^14^. Thus, this paper creates these measures within the HRS sample. This is accomplished by estimating associations between parents and children’s education within the HRS sample between (children’s) birth years of 1920-1960 for each state and decade of birth. This analysis uses a “jackknife procedure” by removing the focal individual from the regression analysis (thus, approximately 20,000 regressions are estimated). The results of these regressions (the parent-child educational association for each state and decade of birth) are then used as the key measure of the social environment in the gene-environment framework below.

Table 1 presents descriptive statistics for the analysis sample. Of the approximately 20,000 HRS respondents with valid parental education reports and state-of-birth information, over 9,000 respondents have genetic data available. Of these, the analysis focuses on the ~7,000 respondents of European ancestry due to a host of issues with applying GWAS coefficients estimated from European samples into non-European samples (see Conley and Fletcher 2017). I also eliminate approximately 500 respondents with birth decade X state mobility measures estimated on fewer than 50 people. Appendix Table 1A presents the summary statistics for the full sample. The average years of schooling completed for “children” (i.e. HRS respondents) is 13.4, while the schooling levels of their fathers (10.14) and mothers (10.63) is lower and more variable. The span of birth years is 1924-1963. While I calculate four versions of state-level mobility that depend on various standardizations of the schooling variables^15^, they are correlated >0.9, so I use output from a years-on-years regression. The polygenic score is standardized by convention and to aid in interpretation in the regression results.

**Table 1.**
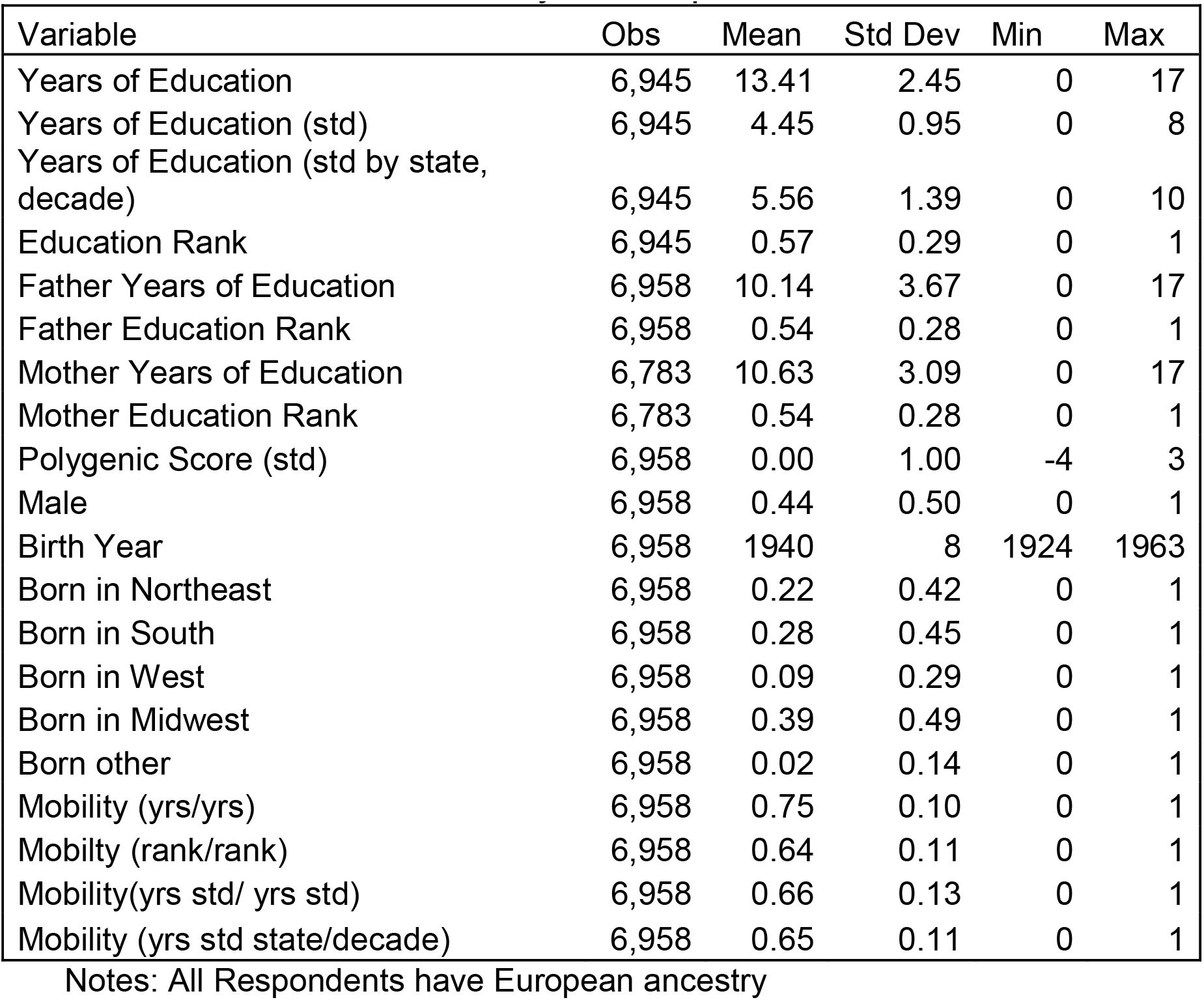
Descriptive Statistics Analysis Sample

## Results

The main empirical strategy takes two steps. The first step is to examine whether there are gene-environment correlations (i.e. confounding) in the data. That is, it is possible that parents with genotypes predicting high education purposefully locate in state with high (or low) educational mobility. Table 2 explores this possibility. While some results are unsurprising—those born in southern states face much lower educational mobility rates than those born in the North East, the result show no evidence that father’s education is correlated with state level mobility nor that child’s polygenic score for education is correlated with state level mobility. Figure 1 presents the densities of children’s polygenic scores stratified by whether the child is born in a high or low (based on a median split of the data) mobility state. Figure 2 benchmarks these results by showing the child’s polygenic score stratified by whether paternal education is high or low (based on a median split of the data). These results suggest the importance of gene-environment correlations for the main analysis is negligible.

**Table 2.**
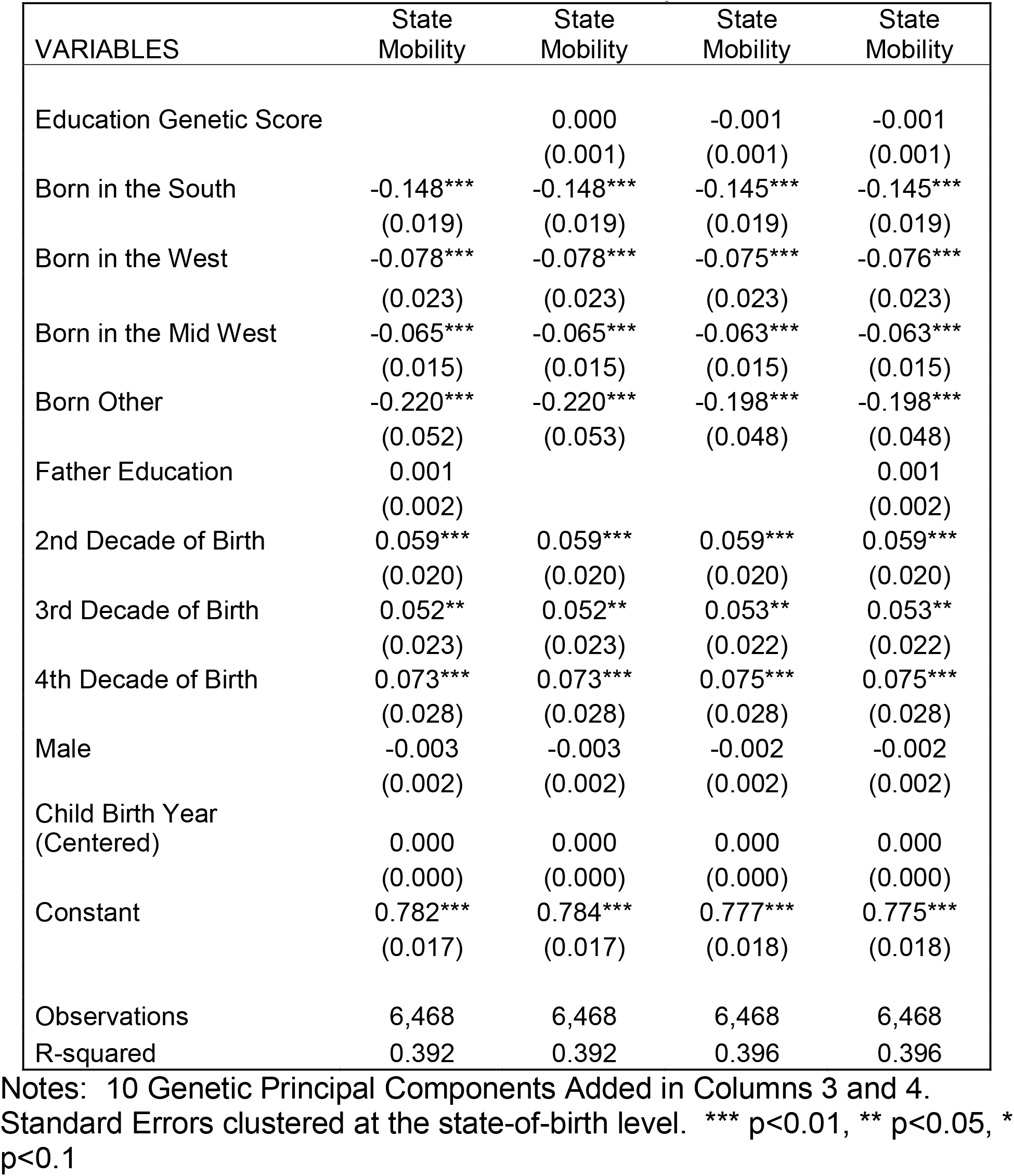
Predictors of State-of-Birth Mobility Measure

| VARIABLES | State<br>Mobility | State<br>Mobility | State<br>Mobility | State<br>Mobility |
| --- | --- | --- | --- | --- |
| Education Genetic Score |  | 0.000<br>(0.001) | -0.001<br>(0.001) | -0.001<br>(0.001) |
| Born in the South | -0.148***<br>(0.019) | -0.148***<br>(0.019) | -0.145***<br>(0.019) | -0.145***<br>(0.019) |
| Born in the West | -0.078***<br>(0.023) | -0.078***<br>(0.023) | -0.075***<br>(0.023) | -0.076***<br>(0.023) |
| Born in the Mid West | -0.065***<br>(0.015) | -0.065***<br>(0.015) | -0.063***<br>(0.015) | -0.063***<br>(0.015) |
| Born Other | -0.220***<br>(0.052) | -0.220***<br>(0.053) | -0.198***<br>(0.048) | -0.198***<br>(0.048) |
| Father Education | 0.001<br>(0.002) |  |  | 0.001<br>(0.002) |
| 2nd Decade of Birth | 0.059***<br>(0.020) | 0.059***<br>(0.020) | 0.059***<br>(0.020) | 0.059***<br>(0.020) |
| 3rd Decade of Birth | 0.052**<br>(0.023) | 0.052**<br>(0.023) | 0.053**<br>(0.022) | 0.053**<br>(0.022) |
| 4th Decade of Birth | 0.073***<br>(0.028) | 0.073***<br>(0.028) | 0.075***<br>(0.028) | 0.075***<br>(0.028) |
| Male | -0.003<br>(0.002) | -0.003<br>(0.002) | -0.002<br>(0.002) | -0.002<br>(0.002) |
| Child Birth Year<br>(Centered) | 0.000<br>(0.000) | 0.000<br>(0.000) | 0.000<br>(0.000) | 0.000<br>(0.000) |
| Constant | 0.782***<br>(0.017) | 0.784***<br>(0.017) | 0.777***<br>(0.018) | 0.775***<br>(0.018) |
| Observations | 6,468 | 6,468 | 6,468 | 6,468 |
| R-squared | 0.392 | 0.392 | 0.396 | 0.396 |

Notes: 10 Genetic Principal Components Added in Columns 3 and 4. Standard Errors clustered at the state-of-birth level. \*\*\* p<0.01, \*\* p<0.05, \* p<0.1

The second analysis step examines the key hypothesis of the paper: I estimate whether the contextual environment may moderate the impacts of genetic endowments on future educational attainments –gene-environment interactions. This analysis is done by interacting the social environmental measure of mobility at the state X time level with the child’s genetic score, while also controlling for main effect of genetics and mobility:

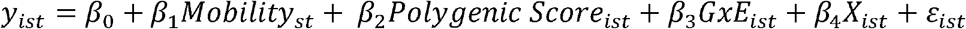

First, Table 3 presents results for the main effects of polygenic score and state mobility without their interaction. The results serve as a benchmark of adjusted versus unadjusted associations between polygenic scores and years of schooling. The first column shows that a one standard deviation increase in the polygenic score is associated with an increase of over 0.6 years of schooling. The second and third column contains estimates of the intergenerational correlation in education between children and fathers (0.26) and children and mothers (0.30) as well as when both measures are included, their sum is 0.35. The final column includes both children’s polygenic score and parental schooling. Comparing column 1 with column 5, it appears that a component of the “genetic effect” could capture enduring “environment” effects of having educated parents.

**Table 3.**
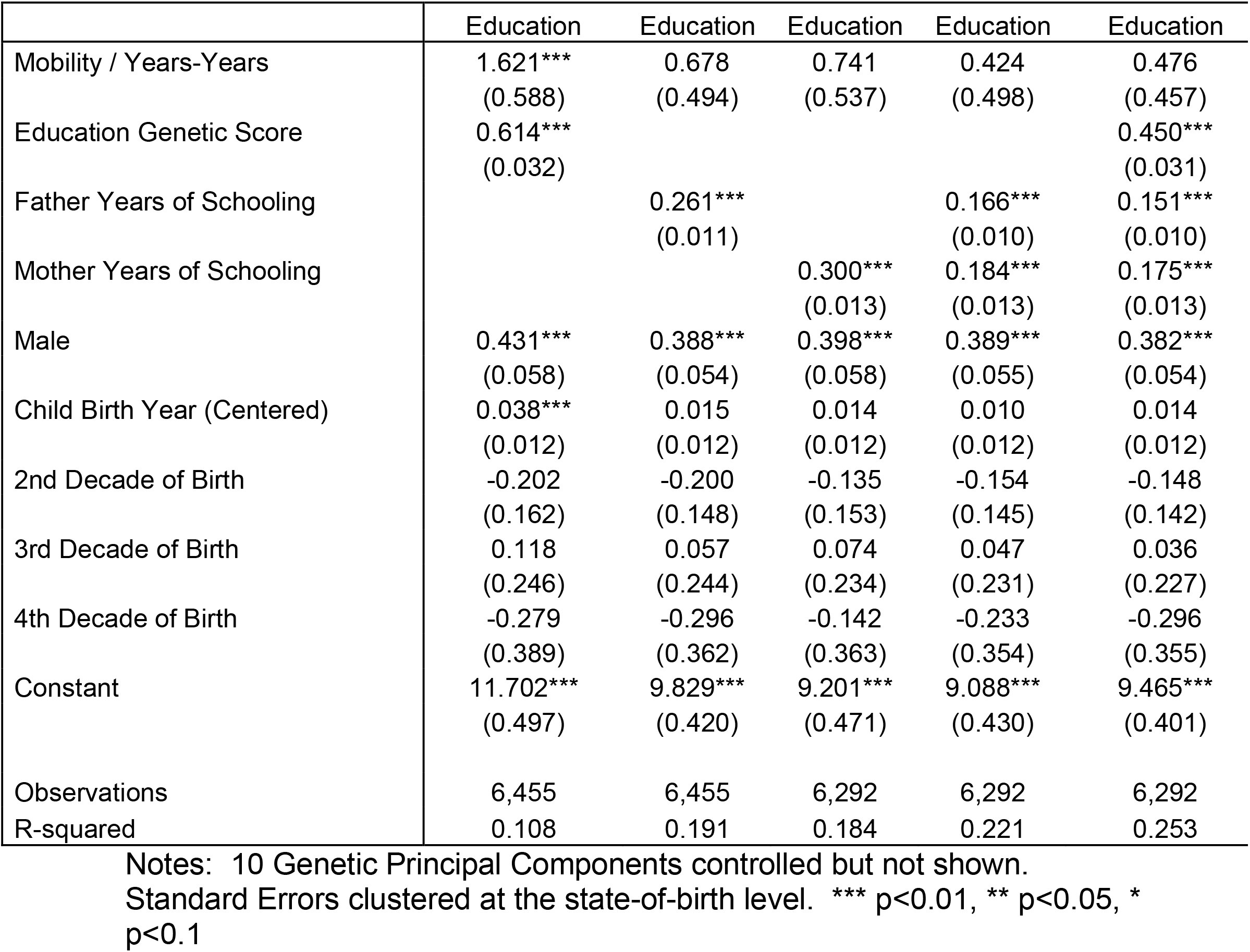
Associations between Parental Education, Genetic Score, and Child Education

Table 4 extends the analysis to include gene-environment interactions. Column 1 repeats results from Table 3. Column 2 shows results that suggest that children born in states with high mobility have lower genetic penetrance in educational attainment than children born in states with low mobility. The results are shown to be robust across measure of mobility in Table 2A in the Appendix.

**Table 4.** Examination of Gene-Environment Interaction between Child Genotype and State Education Mobility: Years of Schooling

|  | Education | Education | Education | Education |
| --- | --- | --- | --- | --- |
| Mobility / Years-Years | 1.621***<br>(0.588) | 1.633***<br>(0.568) | 0.476<br>(0.457) | 0.487<br>(0.445) |
| Education Genetic Score | 0.614***<br>(0.032) | 1.172***<br>(0.232) | 0.450***<br>(0.031) | 0.880***<br>(0.239) |
| Father Years of Schooling |  |  | 0.151***<br>(0.010) | 0.151***<br>(0.010) |
| Mother Years of Schooling |  |  | 0.175***<br>(0.013) | 0.175***<br>(0.013) |
| Mobility X Genetic Score |  | -0.745**<br>(0.316) |  | -0.574*<br>(0.329) |
| Male | 0.431***<br>(0.058) | 0.432***<br>(0.058) | 0.382***<br>(0.054) | 0.383***<br>(0.055) |
| Child Birth Year (Centered) | 0.038***<br>(0.012) | 0.038***<br>(0.012) | 0.014<br>(0.012) | 0.015<br>(0.012) |
| 2nd Decade of Birth | -0.202<br>(0.162) | -0.206<br>(0.162) | -0.148<br>(0.142) | -0.151<br>(0.141) |
| 3rd Decade of Birth | 0.118<br>(0.246) | 0.110<br>(0.245) | 0.036<br>(0.227) | 0.029<br>(0.226) |
| 4th Decade of Birth | -0.279<br>(0.389) | -0.295<br>(0.387) | -0.296<br>(0.355) | -0.309<br>(0.355) |
| Constant | 11.702***<br>(0.497) | 11.707***<br>(0.478) | 9.465***<br>(0.401) | 9.471***<br>(0.389) |
| Observations | 6,455 | 6,455 | 6,292 | 6,292 |
| R-squared | 0.108 | 0.109 | 0.253 | 0.254 |

Notes: 10 Genetic Principal Components controlled but not shown. Standard Errors clustered at the state-of-birth level. \*\*\* p<0.01, \*\* p<0.05, \* p<0.1

The final column of Table 4 adds parental education controls to the analysis. On one hand, adding these parental education controls can help reduce the chances of gene-environment correlations. On the other hand, an analysis that includes measures of parental education (conditioning out the mechanism of genetic transmission from parental genotype to child’s genotype) is difficult to interpret. Indeed, because the genetic transmission is accounted for, the *partial* association between parental educational attainment and children’s educational attainment may best be interpreted as parental investment “effects”; if these proxies are responsive to state mobility rates, then including them in the analysis may be “over controlling” and instead be a mechanism rather than a source of confounding. Again, the results suggest that children born in high mobility states have lower genetic penetrance for polygenic scores associated with educational attainment.

## Conclusion

This paper provides a new look at the transmission of socioeconomic attainments across generations. In particular, a key question in the intergenerational mobility literature is the extent to which genetic transmission of traits from parents to children “lock in” some base level of persistence. In contrast, emerging work on gene-environment interactions suggests the importance of social environments to modulate how genes are expressed, allowing the possibility of “unlocking” intergenerational persistence in socioeconomic attainments. This paper explores this question of gene-environment interactions in the production of educational attainments using the HRS data and conducting an omnibus test of the hypothesis. The analysis uses new measures of genetic endowments, a polygenic score that summarizes genome-wide effects of genetic variation that predicts educational attainment, as well as creates and includes a state and time based estimate of educational mobility to capture broad environmental context during childhood.

The key question is whether this broad measure of the environment interplays with genetic endowments. The key finding is that children who are born in states with high measured educational mobility have lower genetic penetrance in their educational attainments. These findings are unable to adjudicate whether state-level educational mobility, *per se*, or a correlated contextual measure, is the source of the environmental interaction. The results suggest the need to enrich models of attainment and intergenerational mobility to allow for gene-environment interactions and also suggest a need to further study the mechanisms by which contextual measures of mobility decouple children’s genetic endowments from their eventual outcomes.

## Appendix Figures

**Figure 1.**
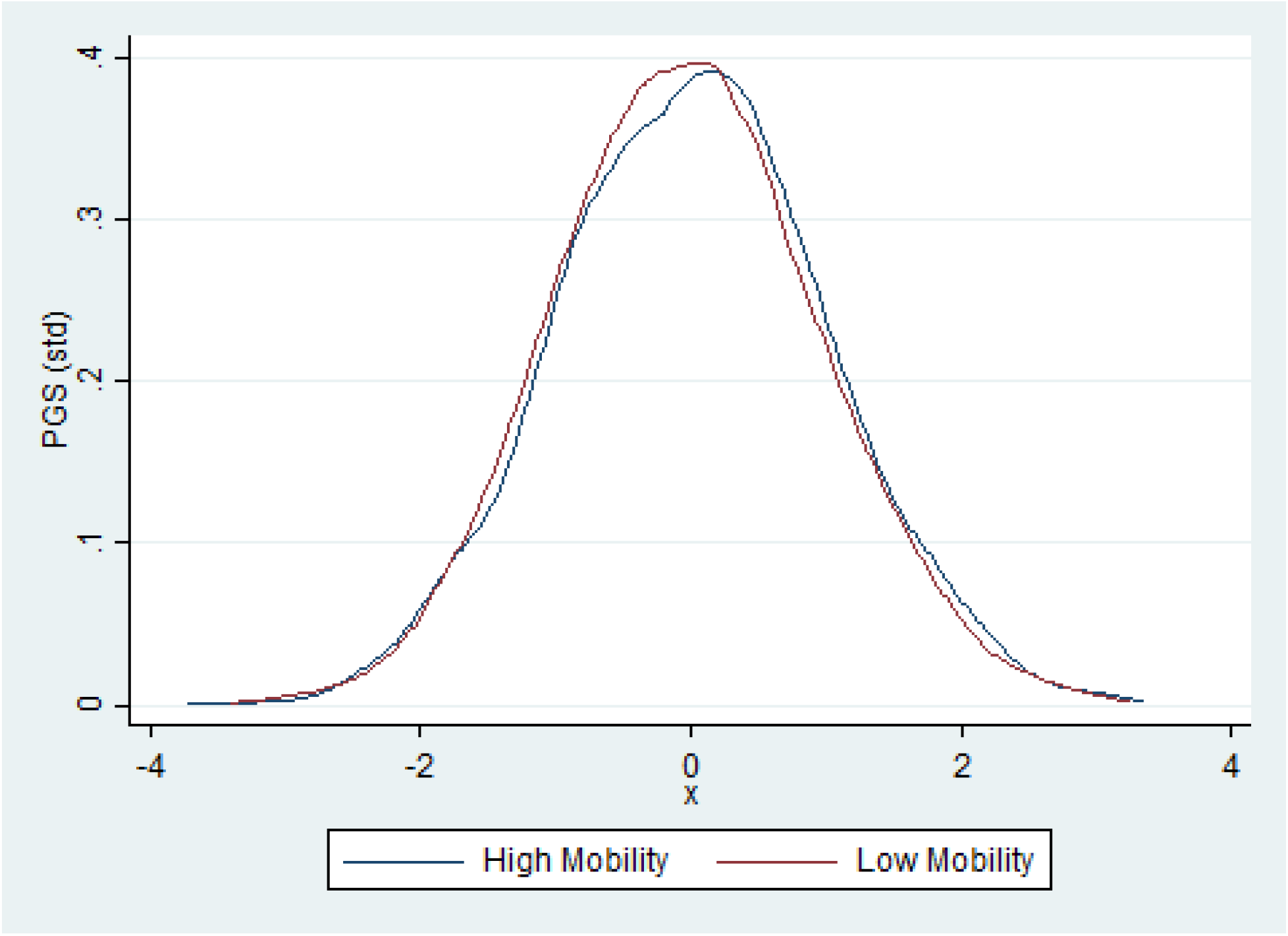
Density of Polygenic Scores Stratified by High vs. Low State Mobility Rates

**Figure 2.**
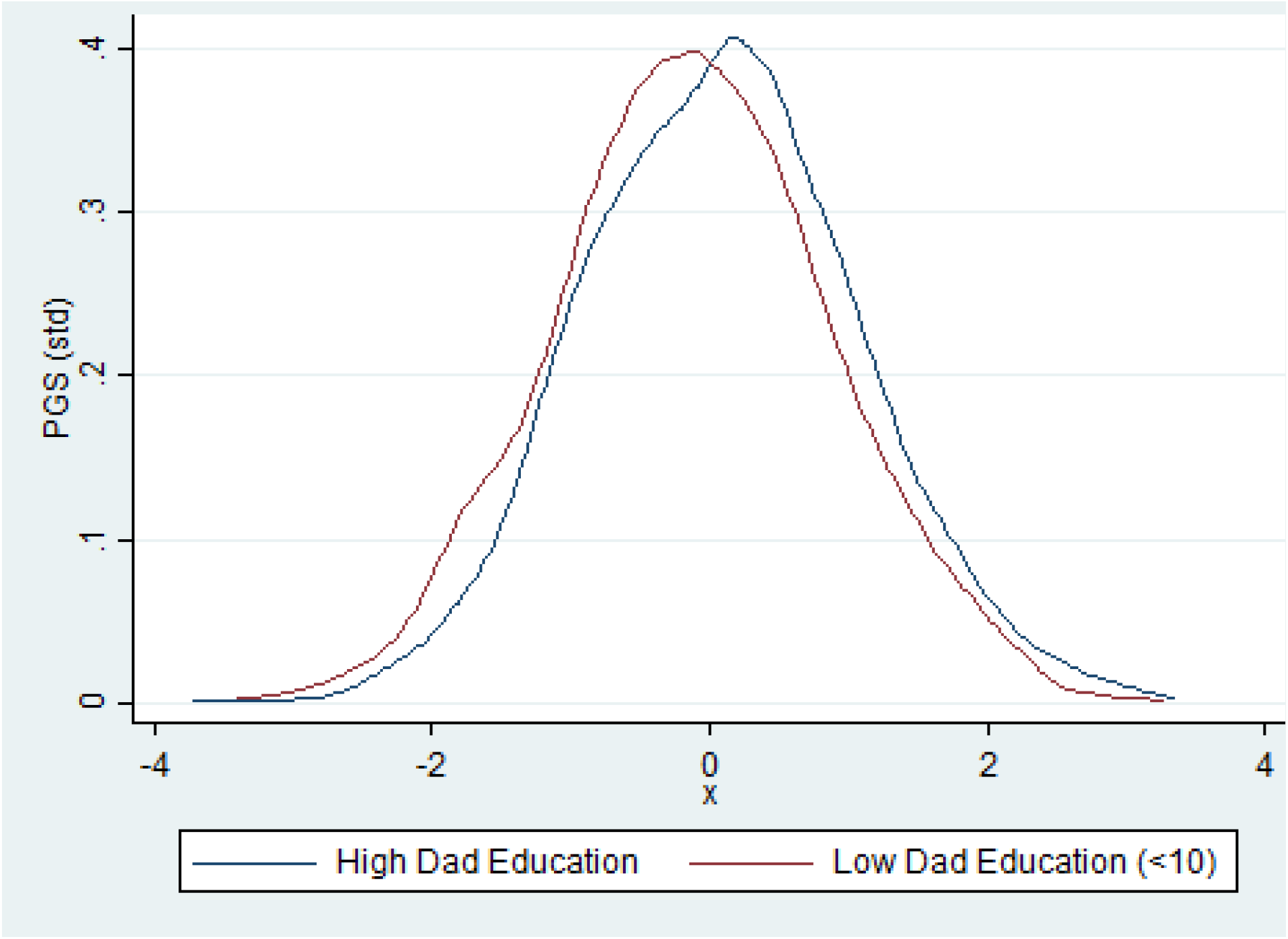
Density of Polygenic Scores Stratified by Paternal Education

## Appendix Tables

**Table 1A.** Descriptive Statistics: Full Sample Compared to Sample with Genotype Data

| Variable | Full Sample |  |  | Analysis Sample |  |  |
| --- | --- | --- | --- | --- | --- | --- |
|  | Obs | Mean | Std. | Obs | Mean | Std. |
| Years of Education | 20,334 | 12.80 | 3.04 | 6,945 | 13.41 | 2.45 |
| Years of Education (std) | 20,334 | 4.40 | 1.26 | 6,945 | 4.45 | 0.95 |
| Years of Education (std by state, decade) | 20,334 | 5.21 | 1.69 | 6,945 | 5.56 | 1.39 |
| Education Rank | 20,334 | 0.53 | 0.30 | 6,945 | 0.57 | 0.29 |
| Log Wealth in 2008 | 11,870 | 11.10 | 4.54 | 6,774 | 11.89 | 3.60 |
| Father Years of Education | 20,385 | 9.49 | 4.13 | 6,958 | 10.14 | 3.67 |
| Father Years of Education (std) | 20,385 | 2.37 | 1.05 | 6,958 | 2.50 | 0.91 |
| Father Years of Education (std by state, decade) | 20,385 | 2.64 | 1.24 | 6,958 | 2.88 | 1.12 |
| Father Education Rank | 20,385 | 0.51 | 0.29 | 6,958 | 0.54 | 0.28 |
| Mother Years of Education | 19,703 | 9.95 | 3.70 | 6,783 | 10.63 | 3.09 |
| Mother Years of Education (std) | 19,703 | 2.79 | 1.09 | 6,783 | 2.91 | 0.86 |
| Mother Years of Education (std by state, decade) | 19,703 | 3.27 | 1.45 | 6,783 | 3.53 | 1.23 |
| Mother Education Rank | 19,703 | 0.51 | 0.28 | 6,783 | 0.54 | 0.28 |
| Polygenic Score (std) | 6,958 | 0.00 | 1.00 | 6,958 | 0.00 | 1.00 |
| Male | 20,385 | 0.46 | 0.50 | 6,958 | 0.44 | 0.50 |
| Birth Year | 20,385 | 1942 | 10 | 6,958 | 1940 | 8 |
| Born in Northeast | 20,385 | 0.19 | 0.39 | 6,958 | 0.22 | 0.42 |
| Born in South | 20,385 | 0.36 | 0.48 | 6,958 | 0.28 | 0.45 |
| Born in West | 20,385 | 0.10 | 0.30 | 6,958 | 0.09 | 0.29 |
| Born in Midwest | 20,385 | 0.29 | 0.45 | 6,958 | 0.39 | 0.49 |
| Born other | 20,385 | 0.06 | 0.25 | 6,958 | 0.02 | 0.14 |
| Mobility (yrs/yrs) | 20,385 | 0.74 | 0.11 | 6,958 | 0.75 | 0.10 |
| Mobility (rank/rank) | 20,385 | 0.62 | 0.11 | 6,958 | 0.64 | 0.11 |
| Mobility(yrs std/ yrs std) | 20,385 | 0.63 | 0.14 | 6,958 | 0.66 | 0.13 |
| Mobility (yrs std state/decade) | 20,385 | 0.63 | 0.12 | 6,958 | 0.65 | 0.11 |

**Table 2A.**
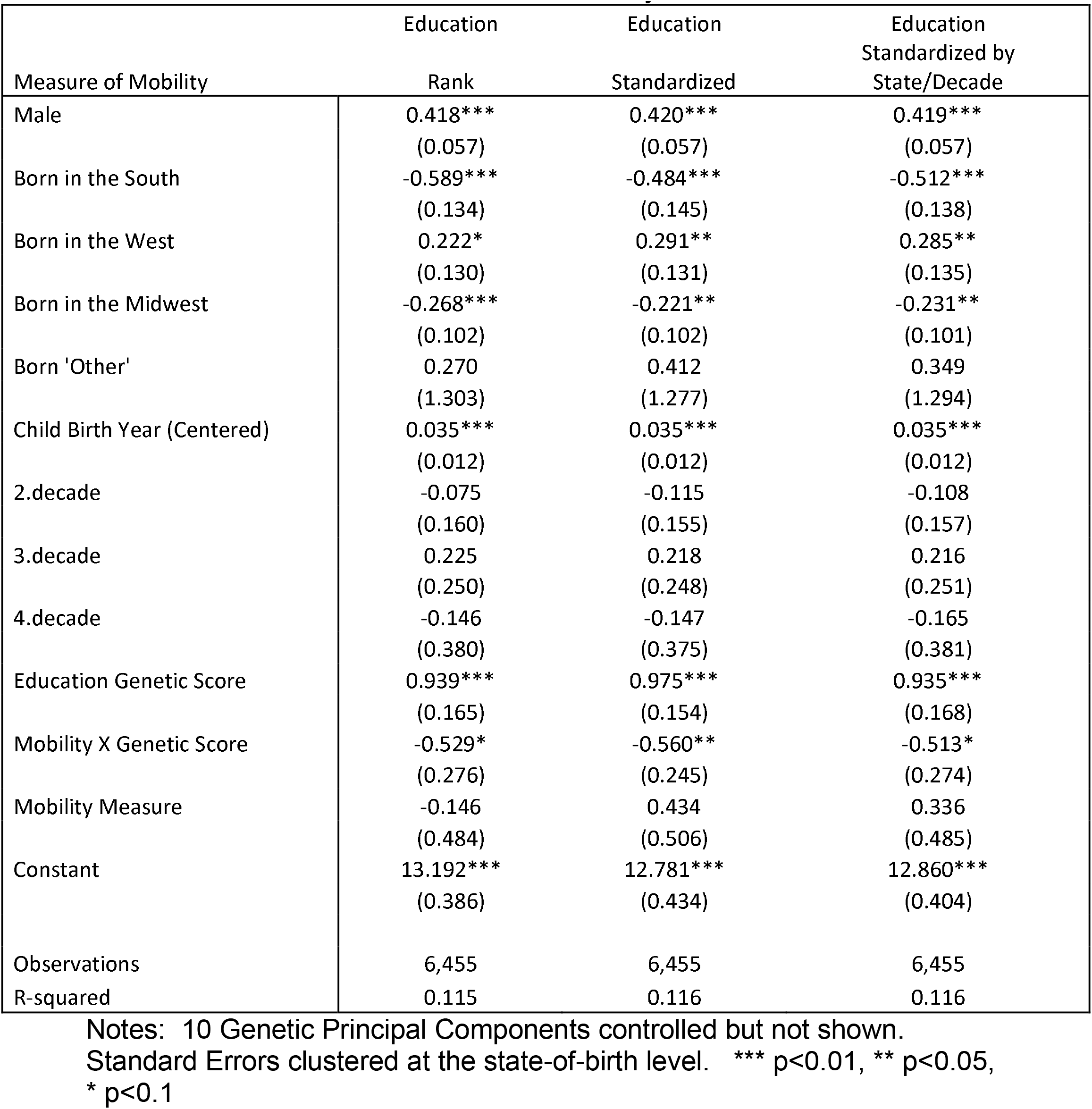
Main Results Across Mobility Measures

## Footnotes

1 The Health and Retirement Study (HRS accession number 0925-0670) is sponsored by the National Institute on Aging (grant numbers NIA U01AG009740, RC2AG036495, and RC4AG039029) and is conducted by the University of Michigan. Additional funding support for genotyping and analysis were provided by NIH/NICHD R01 HD060726. I thank Ben Domingue for help with the polygenic score in HRS and helpful comments from Sandy Black, David Figlio, Eric Grodsky, and participants at the University of Wisconsin Demography Seminar (DemSem), the Ohio State University, University of Georgia, Penn State, Stanford, University of Texas, and the 2017 RC28 Conference and PAA Conference.

2 The typical solution to this issue is to leverage adopted children, which breaks the genetic links between parents and children (e.g. Sacerdote 2007). However, a limitation of this approach is the difficulty of examining both the impacts of environments as well as the potential interactions between genes and environments.

3 For example, studying the differential impacts of school quality based on children’s genetic endowments (gene-environment interaction) is complicated if the parent chooses the school for the child, inducing a gene-environment correlation.

4 The polygenic score, a single scalar measure predicting education, has an R-square for educational attainment in the HRS of over 7%. See also Papageorge and Thom (2016)

5 There has been a recent release of a new polygenic score (https://www.thessgac.org/data) based on a forthcoming paper. Using this new score does not change the main results in the paper.

6 An important paper by Bjorklund et al. (2006) attempted to examine gene-environment interactions by using data on adopted children as well as their biological and adoptive parents. The authors interacted the education levels of the children’s biological (genetic) and adoptive (environmental) parents and found limited effects. Three issues with this approach is that child genotype was inferred from biological parent education, the possibility of selection bias based on the placement of adopted children, and the possibility that adopted children are unrepresentative of all children.

7 Schmitz and Conley (2016) find that (birth date based) eligibility for the Vietnam draft interacts with a genetic score in predicting education attainments of men in the data.

8 Interestingly, the authors did find evidence of interaction between the genetic score of the mother and genetic score of the child in determining the child’s eventual educational attainment. However, without controls for the father’s genotype, it is unclear how to interpret this interaction.

9 See Fletcher and Lehrer (2009) for discussion of this research design.

10 This result is consistent with the conceptual model of Orchids and Dandelions, where some people have genotypes that make the relatively insensitive to environmental variation (dandelions) and others are very sensitive, wilting in bad environments and flourishing in good environments (orchids). See Cook and Fletcher (2015) for evidence.

11 See Domingue et al. (2016) for discussion of mortality selection in the genetic sample.

12 These 2.5 million locations are expanded in a round of imputation to be over 21 million locations. http://hrsonline.isr.umich.edu/index.php?p=xxgen1&_ga=1.238849673.862524756.1380327234

13 Recall, humans are estimated to be over 99.5% genetically identical to one another.

14 Measures of mobility distributed by Raj Chetty begin in 1940 at the state/year level.

15 These include a years-on-years regression, a rank-rank regression where the ranks are calculated over each generation (child/father) separately, a years-on-years regression with the education attainments divided by the standard deviation in the generation following Azam and Bhatt (2015) and a years-on-years regression with the education attainments divided by the standard deviation at the state X decade level with the HRS sample. It may be useful to use external data to standardize the measures, however years of schooling information was first collected in the 1940 census, making it more challenging to standardize the parental education measures.

